# Analyses of Mutational Patterns Induced by Formaldehyde and Acetaldehyde Reveal Similarity to a Common Mutational Signature

**DOI:** 10.1101/2022.03.08.483495

**Authors:** Mahanish J. Thapa, Reena M. Fabros, Salma Alasmar, Kin Chan

## Abstract

Formaldehyde (CH_2_O) and acetaldehyde (C_2_H_4_O) are reactive small molecules produced endogenously in cells as well as being environmental contaminants. Both of these small aldehydes are classified as human carcinogens, since they are known to damage DNA and exposure is linked to cancer incidence. However, the mutagenic properties of formaldehyde and acetaldehyde remain incompletely understood, at least in part because they are relatively weak mutagens. Here, we use a highly sensitive yeast genetic reporter system featuring controlled generation of long single-stranded DNA regions to show that both small aldehydes induced mutational patterns characterized by predominantly C/G → A/T, C/G → T/A, and T/A → C/G substitutions, each in similar proportions. We observed an excess of C/G → A/T transversions when compared to mock-treated controls. Many of these C/G → A/T transversions occurred at T<u>C</u>/<u>G</u>A motifs. Interestingly, the formaldehyde mutational pattern resembles single base substitution (SBS) signature 40 from the Catalog of Somatic Mutations in Cancer (COSMIC). SBS40 is a mutational signature of unknown etiology. We also noted that acetaldehyde treatment caused an excess of deletion events longer than four bases while formaldehyde did not. This latter result could be another distinguishing feature between the mutational patterns of these simple aldehydes. These findings shed new light on the characteristics of two important, commonly occurring mutagens.

## Introduction

Genomic DNA is constantly damaged by intracellular processes (1) and exposure to exogenous damaging agents (2–4). There are many different types of DNA damage. Intracellular DNA damaging processes include, for example: oxidation of nitrogenous bases (5,6); glycosidic bond breakage, which releases a nitrogenous base from its deoxyribose sugar (7–11); single- and double-stranded breaks of the sugar-phosphate backbone (11–13); base alkylation (11,14,15); cytosine deamination to uracil (11,16); and deamination of 5-methylcytosine to thymine (17–19). Examples of exogenous DNA damage include: ultraviolet (UV) light (3); ionizing radiation (4); tobacco (20); aristolochic acid (21); and aflatoxin (22). Mutations are also thought to result from spontaneous ionization or isomerization (i.e., tautomerization) of DNA bases, which can alter base pairing characteristics (23–26).

It is important to note that these processes do not affect all bases equally. Each of the four nitrogenous bases has its own distinct set of chemically reactive moieties (e.g, amines, carbonyls, or labile ring atoms) (27). For any given DNA damaging process or agent, the base(s) with moieties that readily react will be damaged more frequently than bases without such reactive moieties. Local sequence context can also be a key determinant of vulnerability to damage. Mutational signatures are recurrent patterns of base changes that reflect these forms of specificity: the signatures arise naturally because each particular mutagenic process or DNA damaging agent is more likely to affect certain bases in specific contexts more frequently than others (28).

Mutational signatures typically are inferred using a non-negative matrix factorization (NMF) algorithm (29). NMF takes a mutational dataset as input. It initiates by essentially guessing a solution set of constituent signatures with estimated contributions from each putative signature, and then computes the error when attempting to reconstruct the original dataset using that solution set. NMF then tries a slightly different solution set and recomputes the error. This process loops until finding an optimal solution set that stably minimizes reconstruction error. A globally stable solution set is found when different initial conditions all converge to yield that solution set.

NMF analysis can extract reproducible, recurrent patterns of mutations, which often reflect distinct mutagenic processes or DNA damaging agents. There are many mutational signatures with well-established etiologies, including: single base substitution signature 1 (SBS1) from deamination of 5-methylcytosine at <u>C</u>pG motifs; SBS2 and

SBS13 from enzymatic deamination of cytosine at T<u>C</u> motifs by APOBEC deaminases; SBS3 from deficiencies in homologous recombination DNA repair; SBS4 and SBS29 from tobacco smoking and chewing habits, respectively; SBS6, SBS15, SBS21, SBS26, and SBS44 from various deficiencies in DNA mismatch repair; SBS7 from ultraviolet light exposure; SBS10 from mutation of DNA polymerase epsilon; SBS18 from reactive oxygen species; SBS30 and SBS36 from DNA base excision repair deficiencies; and so forth (28). About one-third of currently defined mutational signatures remain of unknown etiology (30).

Previously, the International Agency for Research on Cancer (IARC) named a number of high-priority carcinogens that required further research to fill significant gaps in knowledge (31). Among these high-priority carcinogens are two small aldehyde compounds, formaldehyde (CH_2_O) and acetaldehyde (C_2_H_4_O). Formaldehyde is classified as a known human carcinogen by IARC, based in part on evidence of occupational exposure being associated with nasal and nasopharyngeal cancers (32,33). Formaldehyde is also produced endogenously in cells, as a major metabolic by-product from amino acid metabolism, resulting in high concentrations of up to ∼100 µM in human blood (33). Acetaldehyde is a reactive compound that humans are commonly exposed to as a result of ethanol consumption. Like formaldehyde, acetaldehyde is also classified as a known human carcinogen (34). Alcohol consumption (and the ensuing generation of acetaldehyde by oxidation of ethanol) is associated with higher risk of multiple types of cancer, including: head and neck; esophageal; liver; breast; and colorectal (35).

Understanding the mutagenic characteristics of formaldehyde and acetaldehyde remain important research questions, which can provide valuable insights into the possible roles of these common small aldehydes in cancer mutagenesis and carcinogenesis. Previous attempts to define the mutational patterns induced by formaldehyde and acetaldehyde (e.g., (36,37)) have been rather inconclusive, with no demonstrated link to defined mutational signatures in cancers. Here, we report a more detailed understanding of the mutational characteristics of both formaldehyde and acetaldehyde, and show that the mutational pattern induced by formaldehyde is similar to a common cancer mutational signature that is currently of unknown etiology, namely single base substitution signature 40.

## Materials and Methods

### Reagents and Consumables

Bacto peptone (product code 211677) and yeast extract (212750) were purchased from Becton, Dickinson and Co. (Franklin Lakes, New Jersey). Canavanine (C9758), adenine sulfate dihydrate (AD0028), formaldehyde (F8775), and acetaldehyde (W200344) were purchased from MilliporeSigma (St. Louis, Missouri). Formaldehyde and acetaldehyde solutions were stored in gas-tight tubes in the dark under nitrogen atmosphere. Agar (FB0010), glucose (GB0219), hygromycin (BS725), PCR purification spin column kit (BS654), agarose (D0012), and Tris-Borate-EDTA (TBE) buffer (A0026) were purchased from BioBasic (Markham, Ontario). G418 sulfate (450-130) was purchased from Wisent (St-Bruno, Québec). Q5 PCR kits were purchased from New England Biolabs Canada (Whitby, Ontario). Gas-tight glass tubes with septa (2048-18150) and accessories (2048-11020 and 2048-10020) were purchased from Bellco Glass Inc. (Vineland, New Jersey).

### Yeast Genetics and Mutagenesis

Mutagenesis experiments used the ySR127 yeast strain, a *MATα* haploid bearing the *cdc13-1* temperature sensitive allele. In addition, ySR127 has a cassette of three reporter genes (*CAN1*, *URA3*, and *ADE2*) near the de novo left telomere of chromosome V. These three genes had been deleted from their native loci. Details about ySR127 were described previously (38) and the strain is available upon request.

Formaldehyde mutagenesis experiments were initiated by inoculating single colonies separately into 5 mL of YPDA rich media (2% Bacto peptone, 1% Bacto yeast extract, 2% glucose, supplemented with 0.001% adenine sulfate) in round bottom glass tubes. Cells were grown at permissive temperature (23°C) for three days. Then, cultures were diluted ten-fold into fresh media in gas-tight glass tubes, shifted to restrictive temperature (37°C), and shaken gently at 150 RPM for three hours, with syringe needles inserted through the septa to enable gas exchange. After a three-hour temperature shift, aliquots of formaldehyde stock solution diluted in media were injected into each tube to obtain the reported final concentrations. Samples were then shaken gently at 150 RPM at 37°C for three more hours, in completely sealed gas-tight tubes, to prevent escape of formaldehyde. When formaldehyde treatment was complete, cells were collected by syringe, lightly centrifuged, washed in water, and plated (using a turntable and cell spreader) onto synthetic complete media to assess survival and onto canavanine-containing media with 0.33x adenine to select for mutants (Can^r^ colonies were off-white while Can^r^ Ade^−^ colonies turn red or pink). Care was taken to handle cells gently throughout, as they were quite fragile. Further details of this plating procedure were described in detail previously (39).

Acetaldehyde mutagenesis experiments were carried out similarly. We found that we could simplify the acetaldehyde experiments by using tightly sealed 50 mL polypropylene tubes for the temperature shift and mutagen treatment, presumably because acetaldehyde is less volatile than formaldehyde and does not require as fastidious gas-tight containment. Similar results were obtained for acetaldehyde treatment when using either type of tubes. Statistical analyses and data visualizations were done using base R version 4.1 (40) and tidyverse package version 1.3 (41).

### Illumina Whole Genome Sequencing and Data Analyses

Can^r^ Ade^−^ mutants from formaldehyde and acetaldehyde treatment experiments were collected and reporter gene loss of function phenotypes were verified as described previously (38). Briefly, Can^r^ red/pink mutants were streaked on YPDA plates. A single colony from each streak was patched onto YPDA. Patches were then replica plated onto glycerol, adenine dropout, canavanine, and uracil dropout media. Mutants that grew on glycerol (i.e., were respiration competent), and were Can^r^ Ade^−^ Ura^+^ were considered suitable for sequencing. Can^r^ Ade^−^ Ura^−^ mutants were avoided because those isolates sometimes turn out to be telomere truncations. Mutants from 4, 6, and 8 mM formaldehyde exposure were chosen for sequencing as these had similar induced mutation frequencies. For acetaldehyde, mutants from 75 mM treatment were selected for sequencing, as this concentration was most mutagenic.

Illumina library preparation and WGS were outsourced to Genome Québec (McGill University, Montréal) or performed on an Illumina MiSeq in our lab. Sequencing reads were uploaded to the NCBI SRA (National Center for Biotechnology Information Sequence Read Archive), accessions PRJNA839792 and PRJNA574140. Bowtie2 version 2.3.5.1 (42), SAMtools 1.9 (43), and bcftools 1.9 (44) were used to map the Illumina reads and call variants. Variants with quality score < 30 and/or with sequencing coverage < 10 were filtered out. The resulting VCF files were passed to MutationalPatterns version 3.6.3 (45) for further analysis and visualization. Other numerical and statistical analyses, and data visualizations were done using base R version 4.1 (40) and tidyverse package version 1.3 (41).

For trinucleotide frequency correction, the Biostrings package version 2.38.0 (46) was used to extract trinucleotide counts for the ySR127 yeast and mm10 mouse reference genomes. Following the convention for reporting mutational signatures, counts for each trinucleotide motif centered on C or T were summed with the counts of their respective reverse complements. The proportion of each trinucleotide was then calculated. To infer the expected pattern in mouse, the frequency of each of the 96 channels of a yeast mutational pattern was multiplied by the ratio of corresponding trinucleotide proportions in mouse vs. in yeast. For example, if a given trinucleotide motif is half as abundant in mouse as in yeast, the corresponding expected frequency of mutations in mouse would be scaled by a factor of 0.5 relative to the observed frequency in yeast data.

## Results

### Formaldehyde- and acetaldehyde-induced mutagenesis

We began by assessing mutagenesis and toxicity induced by addition of formaldehyde or acetaldehyde. These experiments were done using a haploid yeast strain (ySR127) that forms long regions of sub-telomeric single-stranded DNA (ssDNA) when shifted to 37°C due to the *cdc13-1* temperature sensitive point mutation (47). At 37°C the cdc13-1 protein dissociates from telomeres, triggering enzymatic resection of unprotected chromosome ends, which in turn activates the DNA damage checkpoint to arrest cells in G_2_ (47). The reporter genes *CAN1*, *ADE2*, and *URA3* had been deleted from their native loci and reintroduced to the left sub-telomeric region of chromosome V (38). This mutagenesis system is very well suited to studying weak mutagens, as ssDNA is more prone to mutation than double-stranded DNA and repair using the complementary strand is not possible. This latter point is an important consideration, since DNA lesions induced by formaldehyde in duplex DNA are potential substrates for nucleotide excision repair (48). The ssDNA system was used previously to study the mutagenic properties of bisulfite and human APOBEC3G cytidine deaminase (38); abasic sites (49); reactive oxygen species (50); human APOBEC3A and APOBEC3B cytidine deaminases (51); and alkylating agents (52).

We treated temperature-shifted cells with increasing concentrations of formaldehyde or acetaldehyde. Care was taken to seal the formaldehyde-treated samples in gas-tight tubes; otherwise, the formaldehyde would simply volatilize into the gaseous phase and escape into the atmosphere. Increasing concentrations of formaldehyde resulted in lower viability (see Figure 1a). While lower concentrations are relatively well tolerated, 8 mM formaldehyde reduced viability below 50%. Formaldehyde-induced inactivation of *CAN1* was detected from as little as 2 mM treatment (median gene inactivation frequency of 3.3 × 10^−4^, see Figure 1b). Mutagenesis plateaued from 4 to 8 mM formaldehyde, with median mutation frequencies of ∼1.5 × 10^−3^. Mock treated cells (i.e., 0 mM formaldehyde) had median mutation frequency of only 1.2 × 10^−4^. These results show that when the experiments are set up properly to contain the mutagen, formaldehyde is clearly mutagenic to our ssDNA model system.

**Figure 1:**
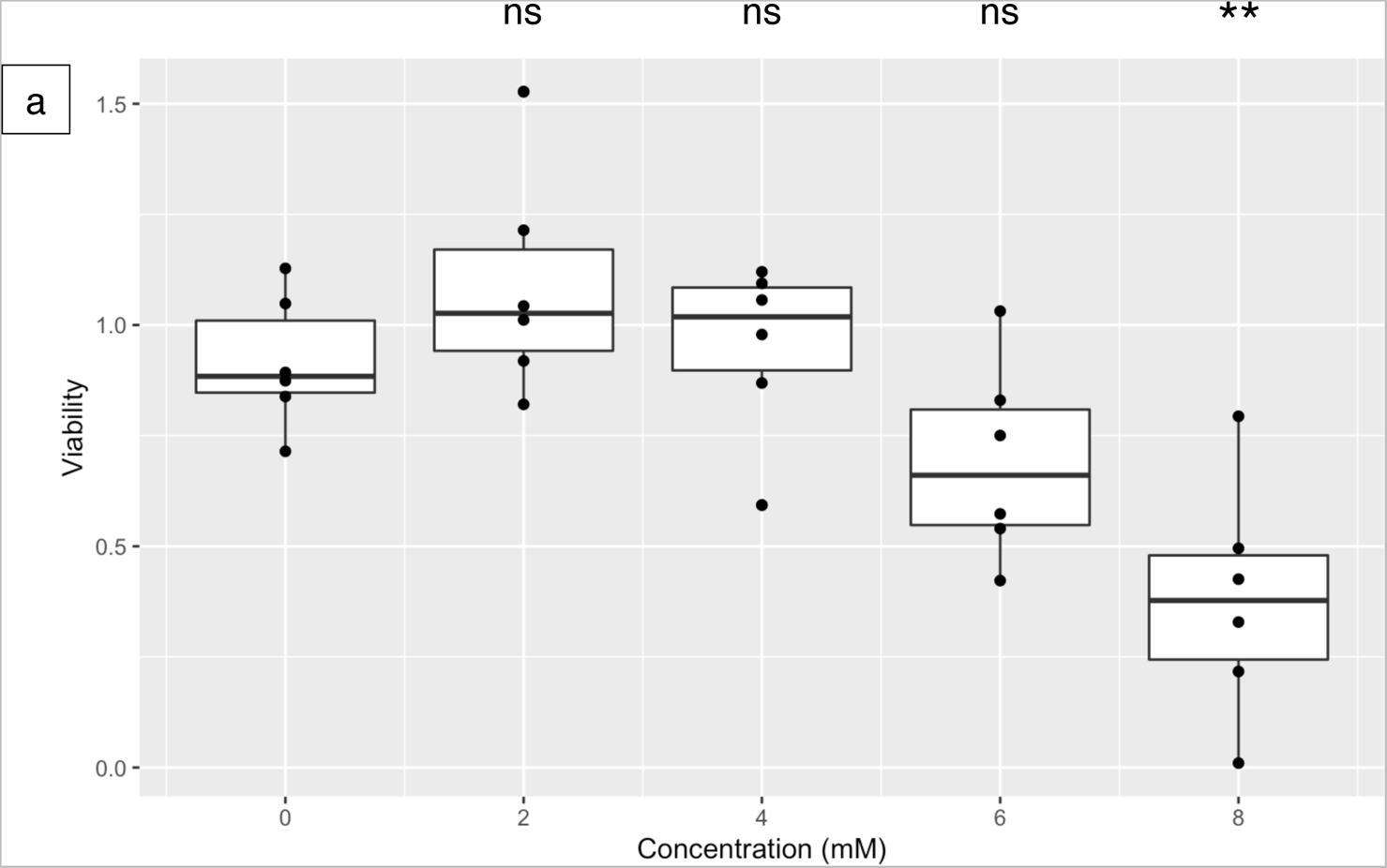
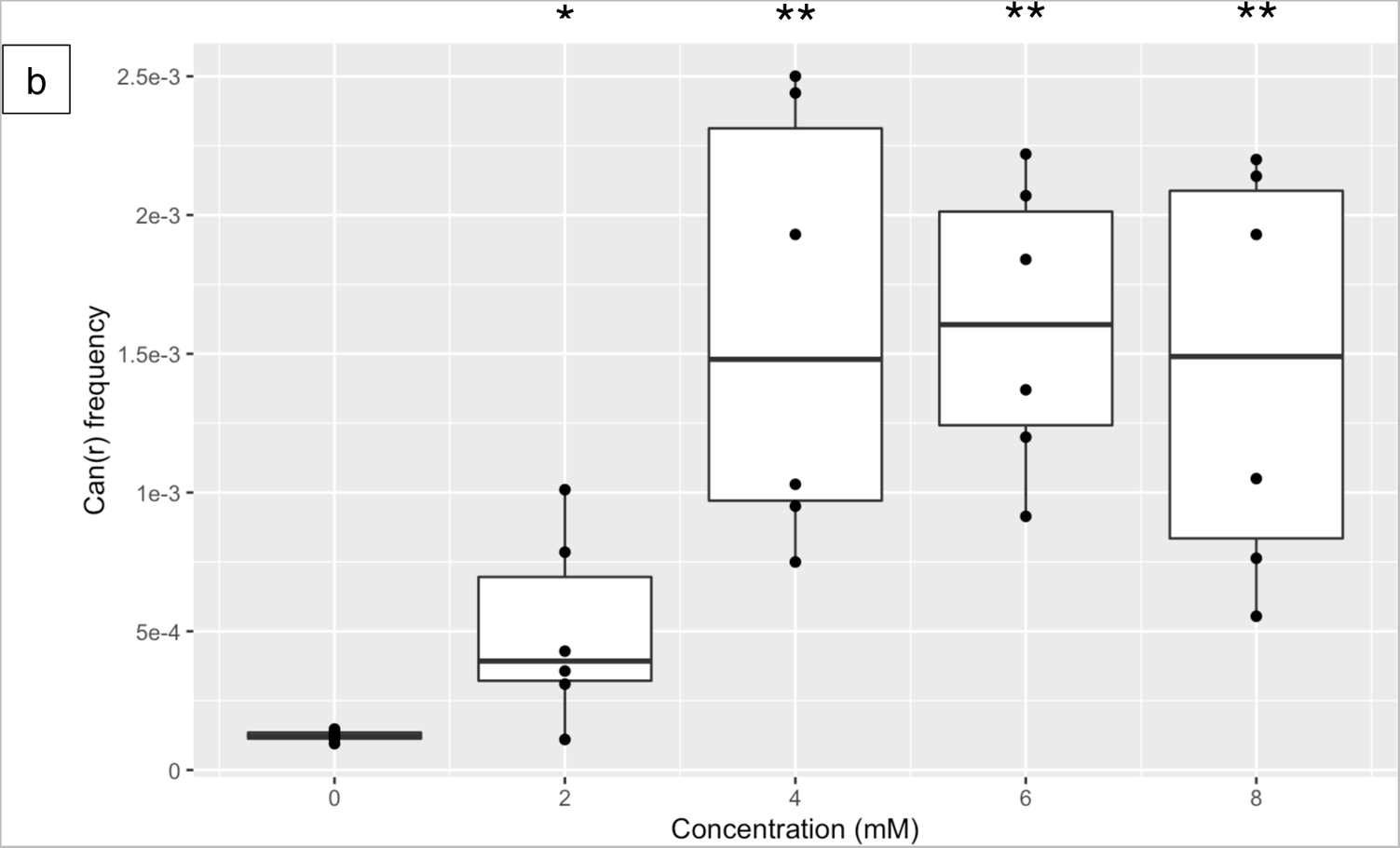
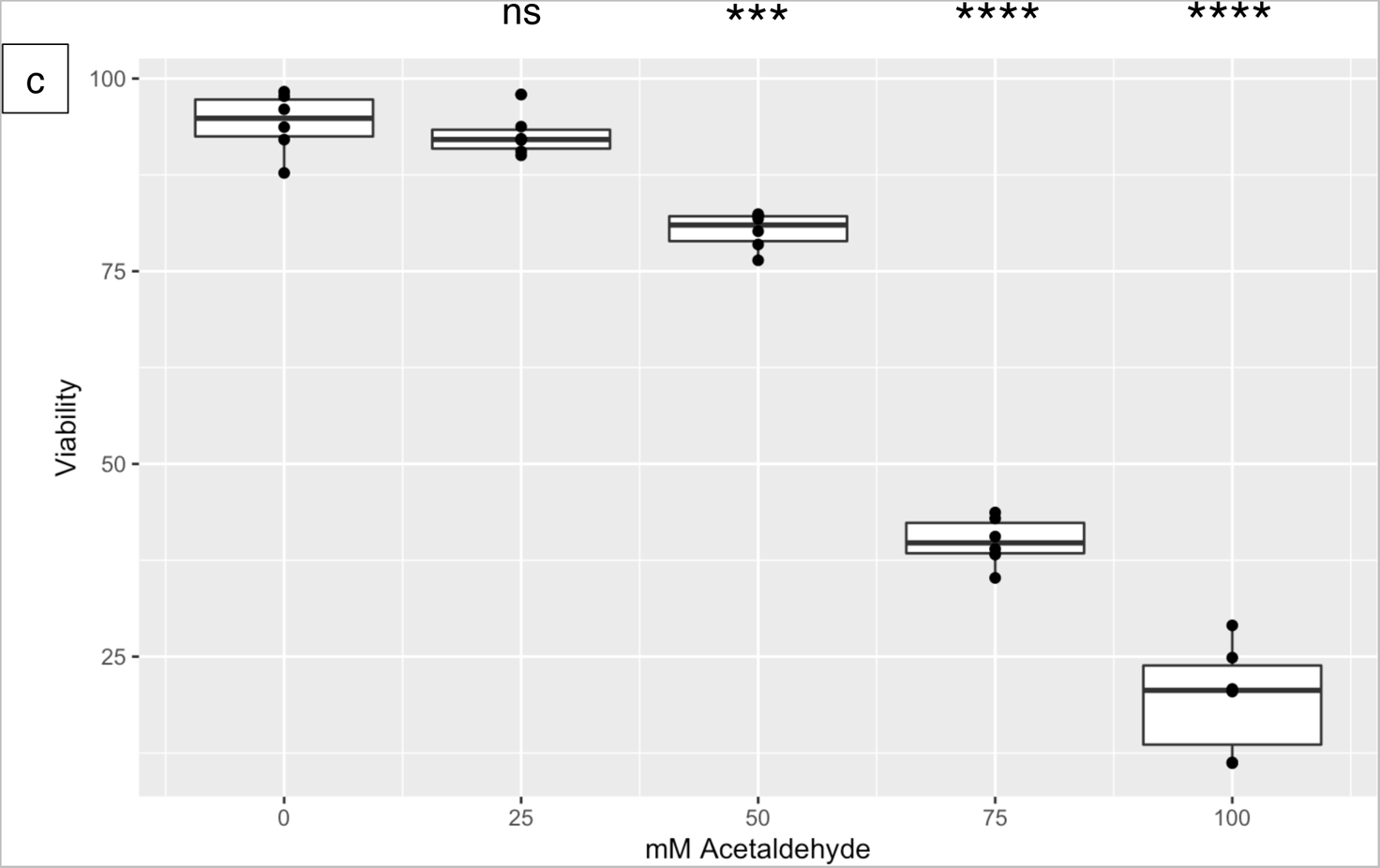
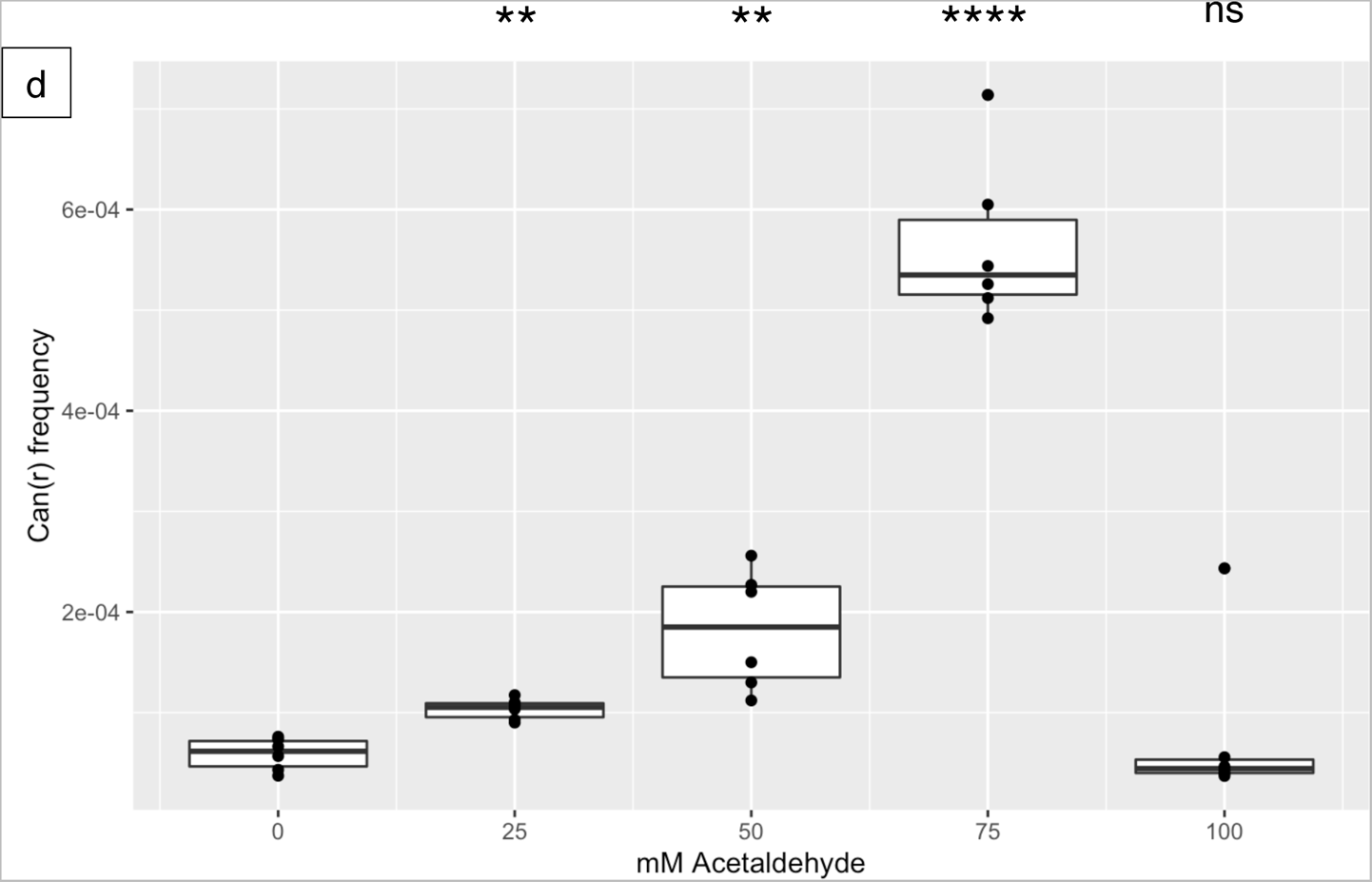
(a) Viability and (b) *CAN1* inactivation frequency of yeast treated with 0, 2, 4, 6, or 8 mM formaldehyde. (c) Viability and (d) *CAN1* inactivation frequency of yeast treated with 0, 25, 50, 75, or 100 mM acetaldehyde. Data are from six biological replicates for each aldehyde. * denotes P < 0.05, ** denotes P < 0.01, *** denotes P < 0.001, **** denotes P < 0.0001, and ns denotes no significant difference by paired t-test.

Cells were considerably more tolerant of higher concentrations of acetaldehyde. We tested concentrations from 25 to 100 mM. Cells treated with lower concentrations (25 and 50 mM) retained high viability, but higher concentrations induced significant lethality (see Figure 1c). Unlike formaldehyde, the mutagenesis induced by acetaldehyde did not show a plateau. Instead, here was a gradual increase in *CAN1* inactivation frequency when treated with 25 and 50 mM acetaldehyde (see Figure 1d). Mutation frequency peaked at over 5 × 10^−4^ when cells were treated with 75 mM acetaldehyde.

Interestingly, treatment with 100 mM acetaldehyde did not result in detectable mutagenesis while viability was reduced to below 25%. This suggests that the cells which sustained high levels of DNA damage by 100 mM acetaldehyde likely suffered considerable cytotoxic damage as well and did not survive.

### Formaldehyde and acetaldehyde both induce an excess of C/G > A/T transversions

We collected mutagenized isolates for Illumina whole genome sequencing to determine what kinds of genetic variants were induced by either formaldehyde or acetaldehyde treatment. The genomes mutagenized by either small aldehyde were compared to control genomes that were not treated by either. Analysis of the untreated control genomes revealed a mutational pattern where C/G > T/A and T/A > C/G transitions outnumbered the four types of transversions (namely C/G > A/T, C/G > G/C, T/A > A/T, and T/A > G/C, see Figure 2a), similar to what we had observed previously (53). By comparison, formaldehyde and acetaldehyde treatment both caused a relative increase of C/G > A/T transversions (see Figure 2b and 2c). While these substitutions accounted for 11% of the mutational spectrum in untreated controls, this fraction rose to about 17% in the aldehyde-mutagenized genomes. This increase is a common characteristic of mutagenesis caused by small aldehydes in regions of single-stranded DNA.

**Figure 2:**
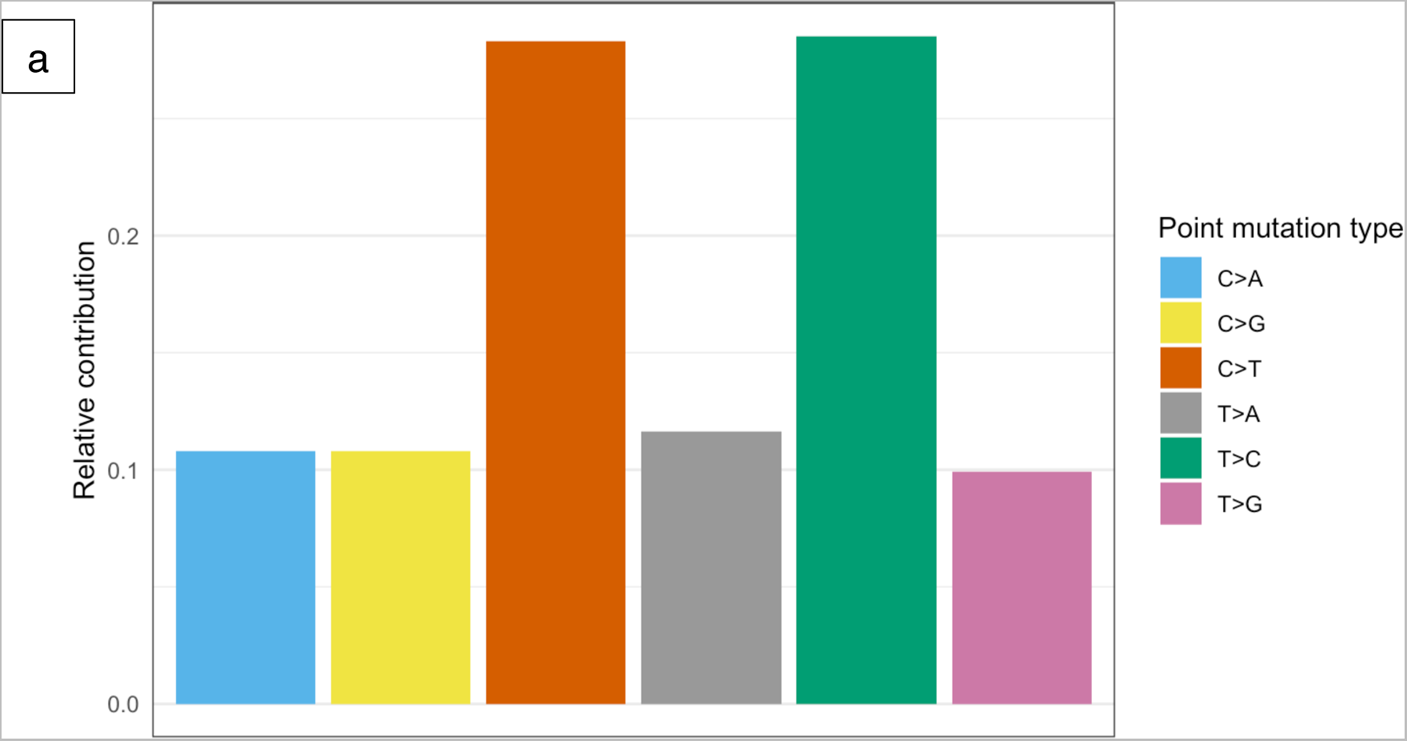
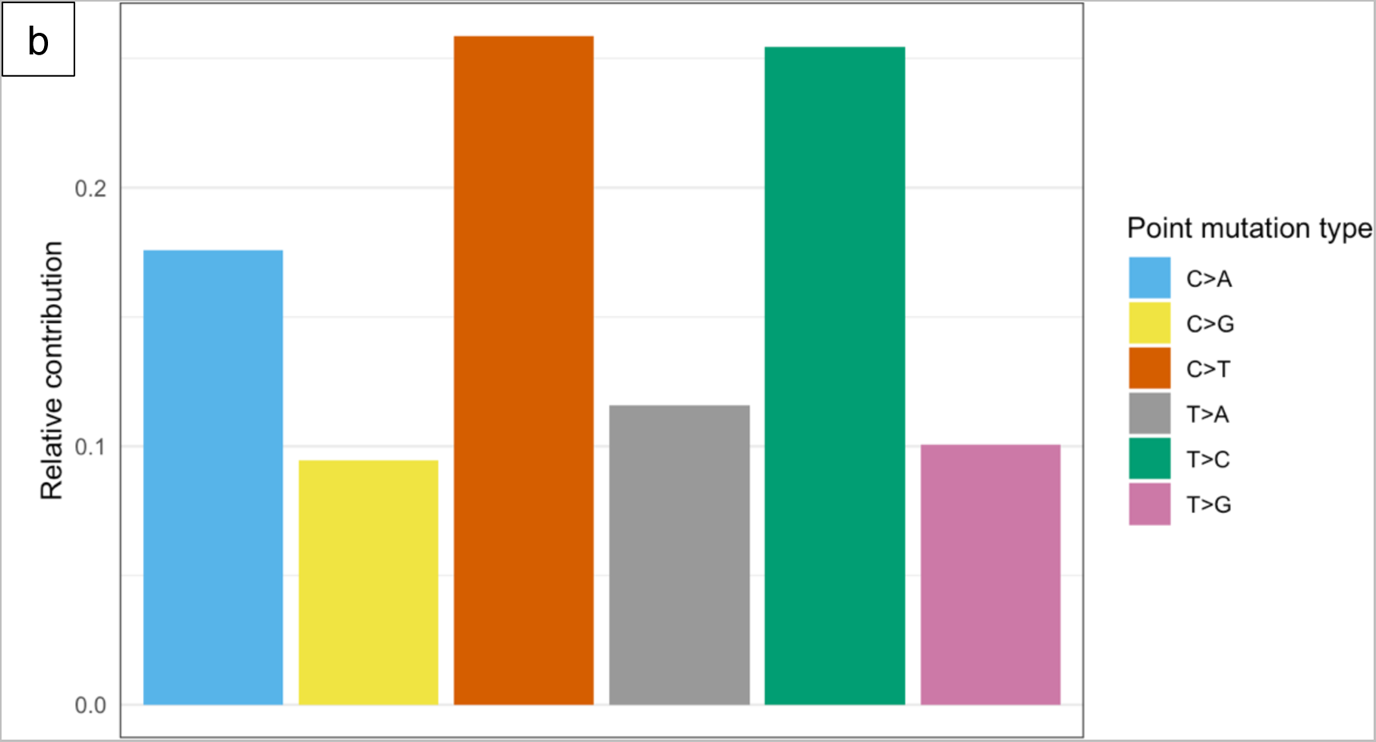
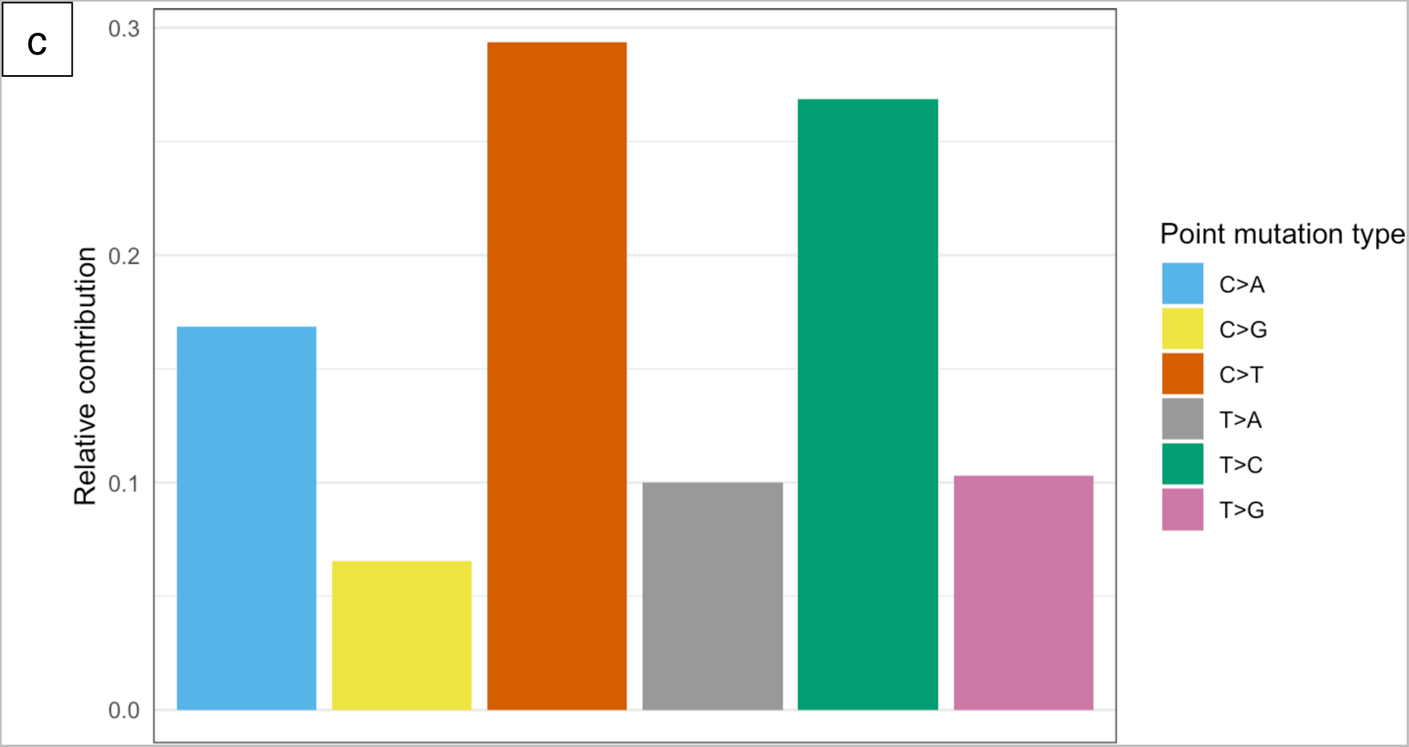
Base substitution types for (a) untreated controls, (b) formaldehyde, and (c) acetaldehyde. Treatment with either aldehyde caused a higher proportion of C/G > A/T transversions.

### Acetaldehyde induces deletions of five or more bases, but formaldehyde does not

We also analyzed short insertions and deletions (indels) to determine if treatment with either small aldehyde can induce these genetic changes. The profile of short indels in untreated controls consists mainly of insertions of five or more bases, with smaller proportions of shorter insertions as well as deletions of five or more bases (see Figure 3a). The profile of short indels in formaldehyde-mutated genomes is essentially the same as in untreated control genomes, i.e., we did not find evidence that formaldehyde induces a higher proportion of any type of indels (see Figure 3b). In contrast, there was a notable difference in the acetaldehyde-induced profile of indels: an excess of deletions of five or more bases was observed (24% in acetaldehyde vs. 12% in controls, see Figure 3c). This is a distinguishing property of acetaldehyde-induced DNA damage in the ssDNA system.

**Figure 3:**
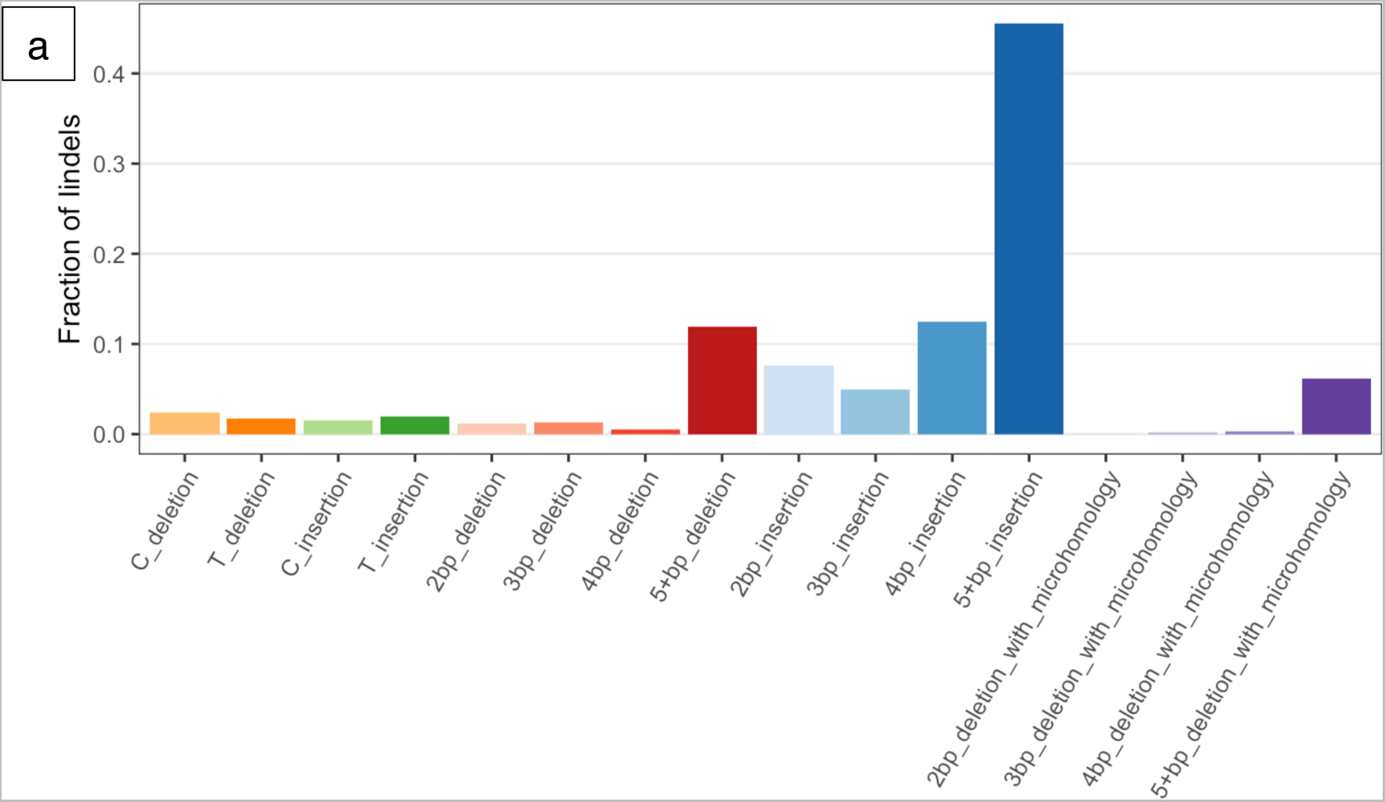
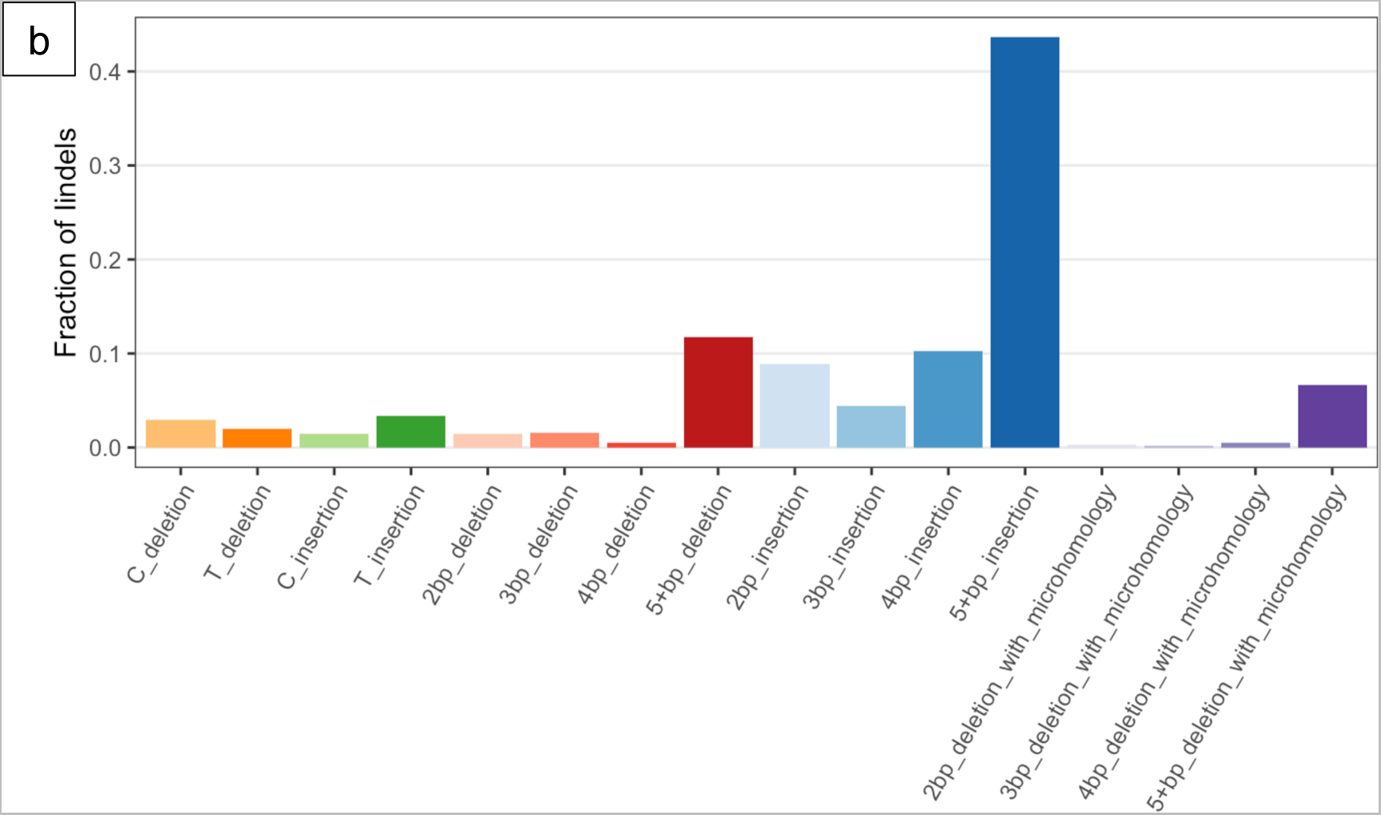
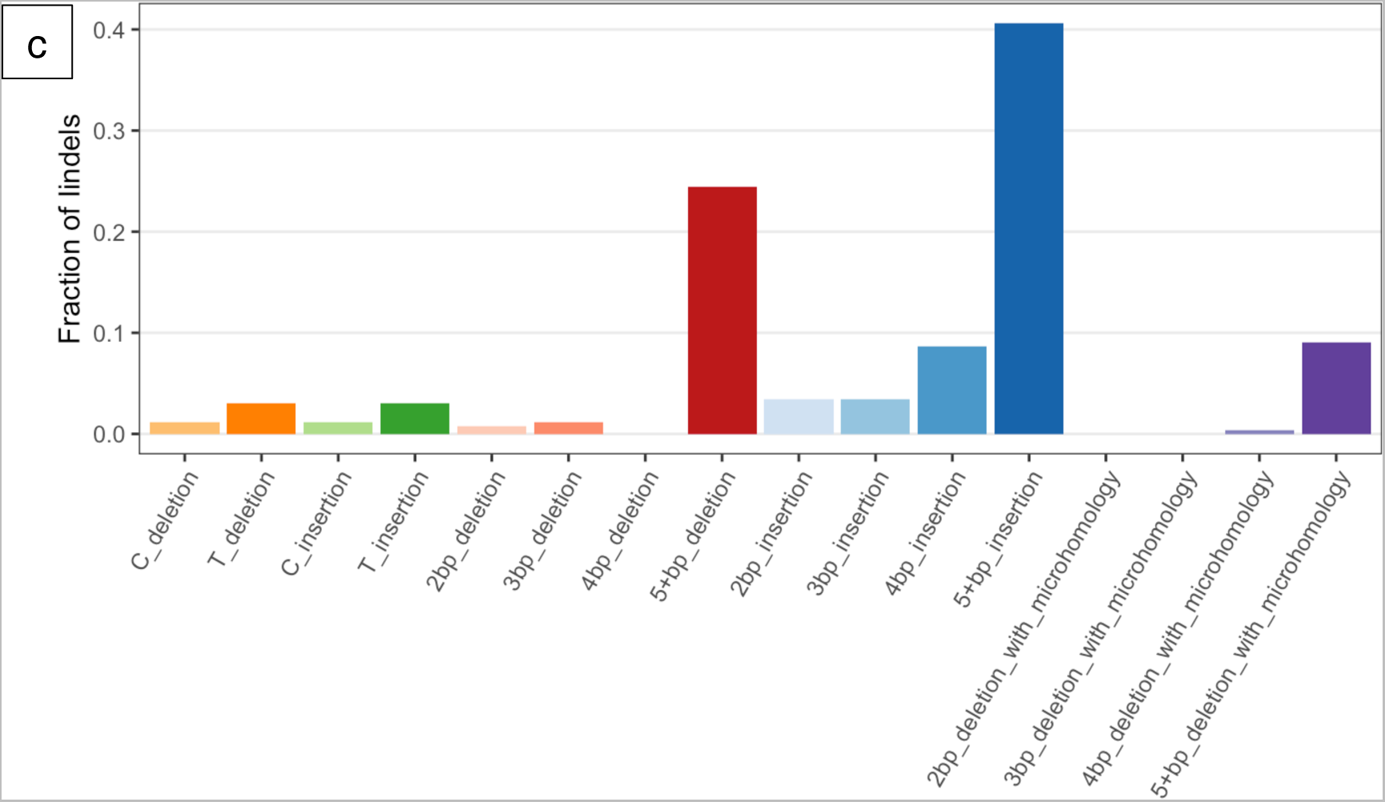
Small indels from (a) untreated controls, (b) formaldehyde, and (c) acetaldehyde. The different categories comprise: single base deletions or insertions at C/G or T/A base pairs; 2, 3, 4, or 5+ base pair deletions or insertions; and 2, 3, 4, or 5+ base pair deletions with microhomology at break points. Acetaldehyde treatment induces an increased proportion of 5+ base pair deletions (without microhomology).

### Formaldehyde and acetaldehyde produce distinct mutational patterns

To investigate the mutational properties of the small aldehydes in more detail, we plotted their mutational profiles in the 96-channel format of the COSMIC mutational signatures. By this convention, all substitutions are reported as originating from a pyrimidine base, i.e., same as the mutation spectra reported above. In addition, the 96-channel profiling features trinucleotide motifs consisting of the mutated base, flanked by an adjacent base 5ʹ and 3ʹ. Cosine similarity is a metric for comparing mutational patterns, yielding a maximum value of exactly 1 for two identical patterns (29). The mutational pattern of formaldehyde is similar to untreated controls (cosine similarity = 0.93), but the excess of C/G > A/T transversions is nonetheless evident (see Figure 4a). The mutational pattern of acetaldehyde is more dissimilar vs. the profile of untreated controls (cosine similarity = 0.868), but again with a noticeable excess of C/G > A/T substitutions (see Figure 4b). When comparing the formaldehyde and acetaldehyde profiles directly to one another, the cosine similarity value is 0.882, showing some similarities but also clear differences in the C/G > T/A channels especially (see Figure 4c).

**Figure 4:**
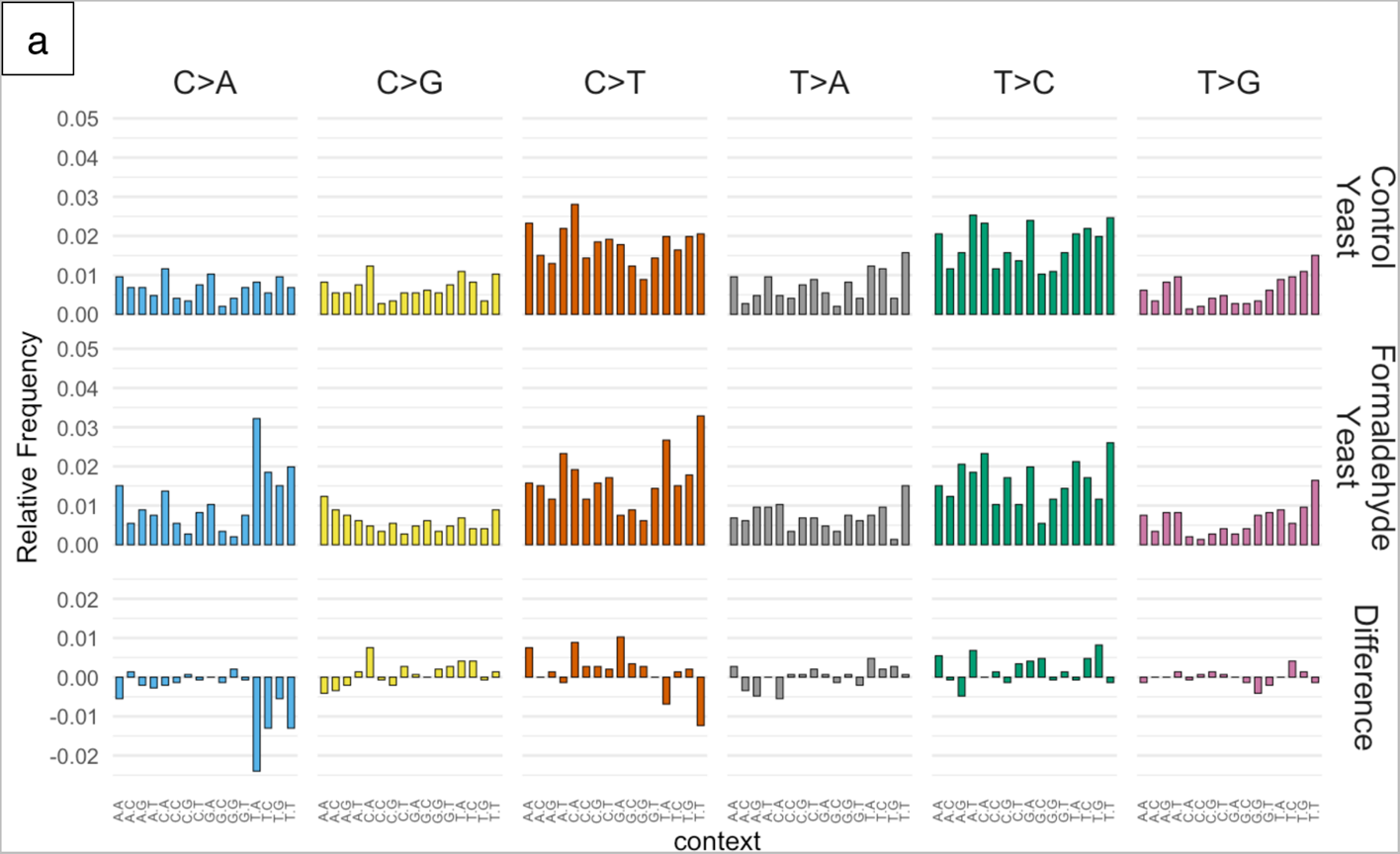
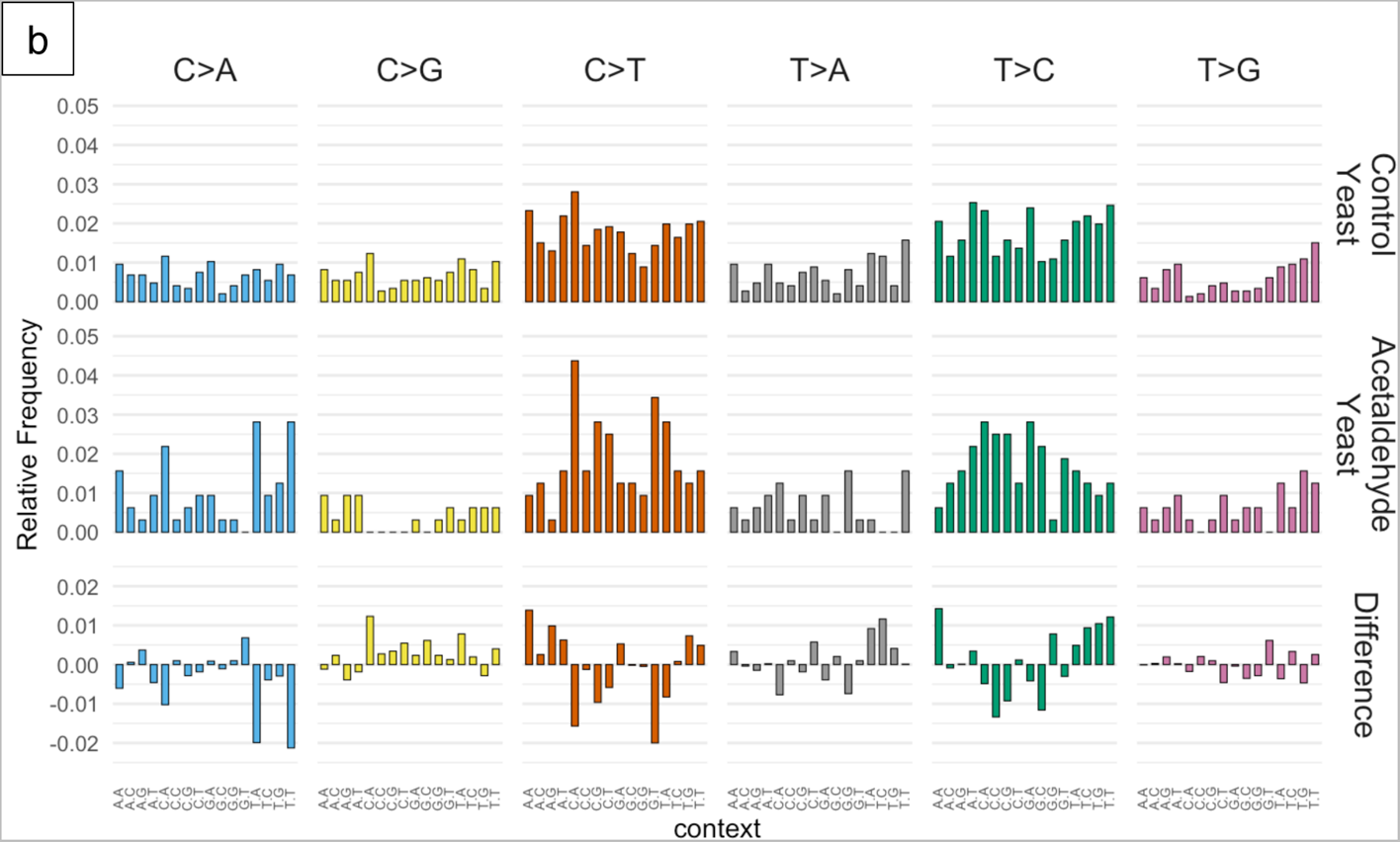
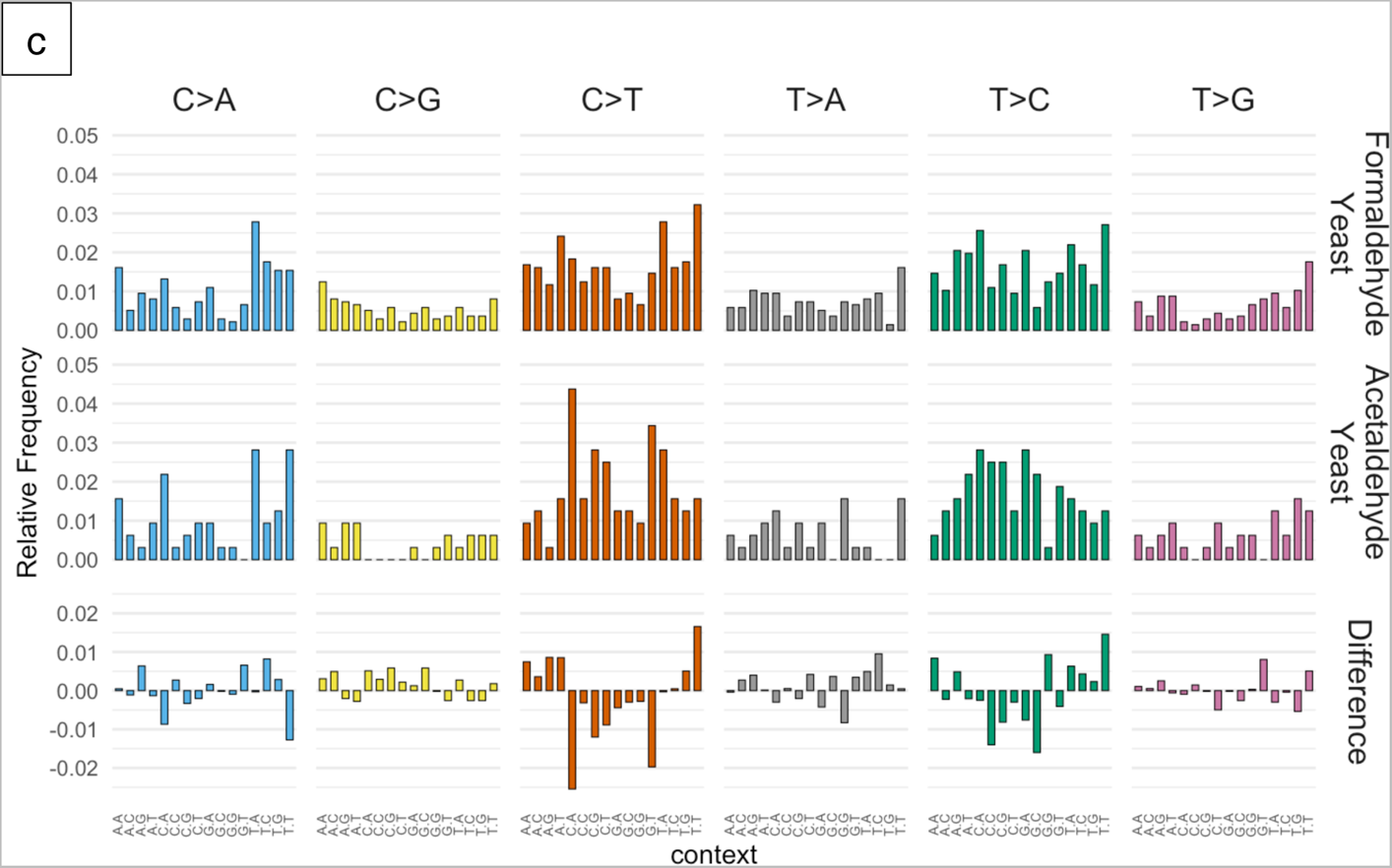
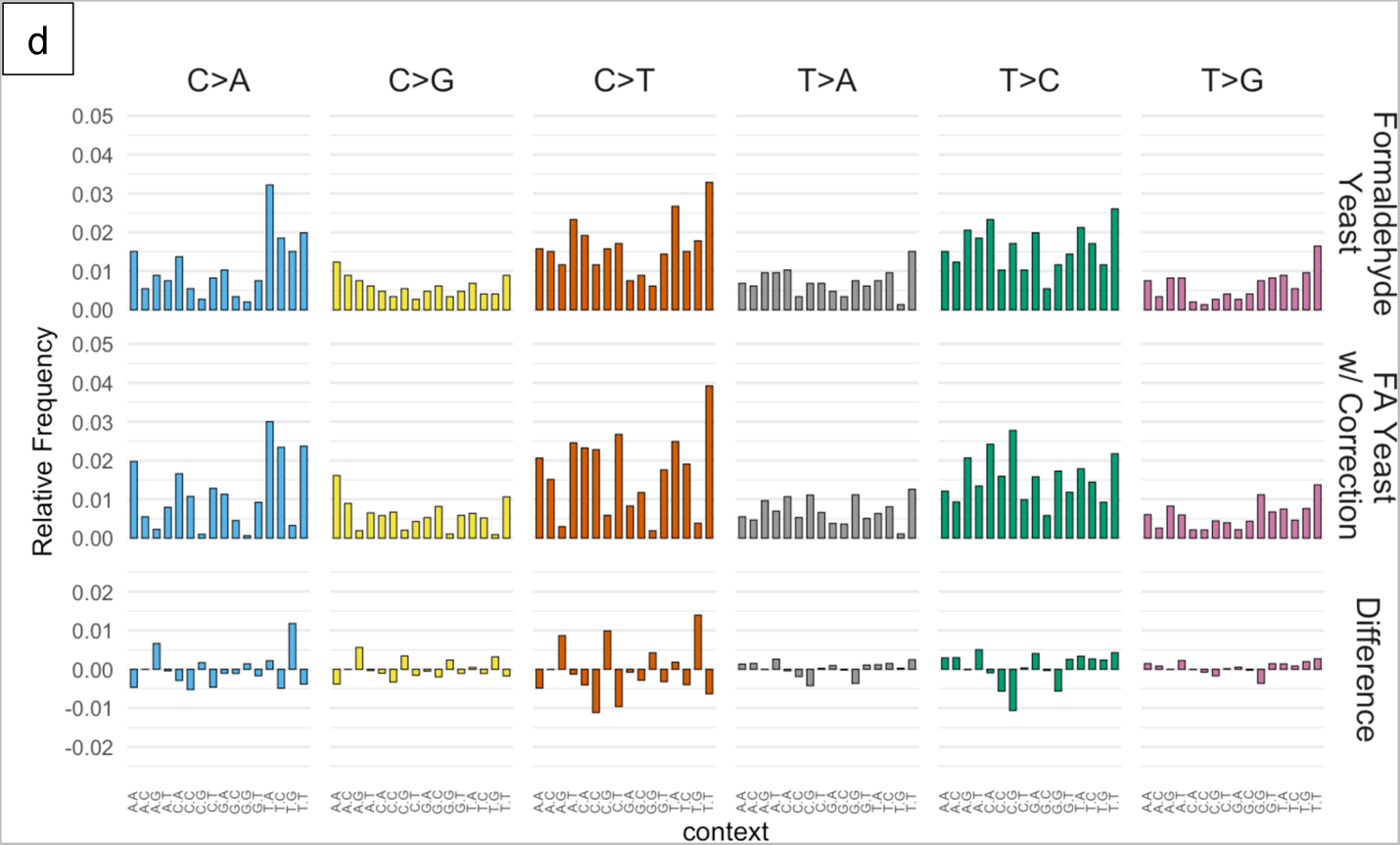
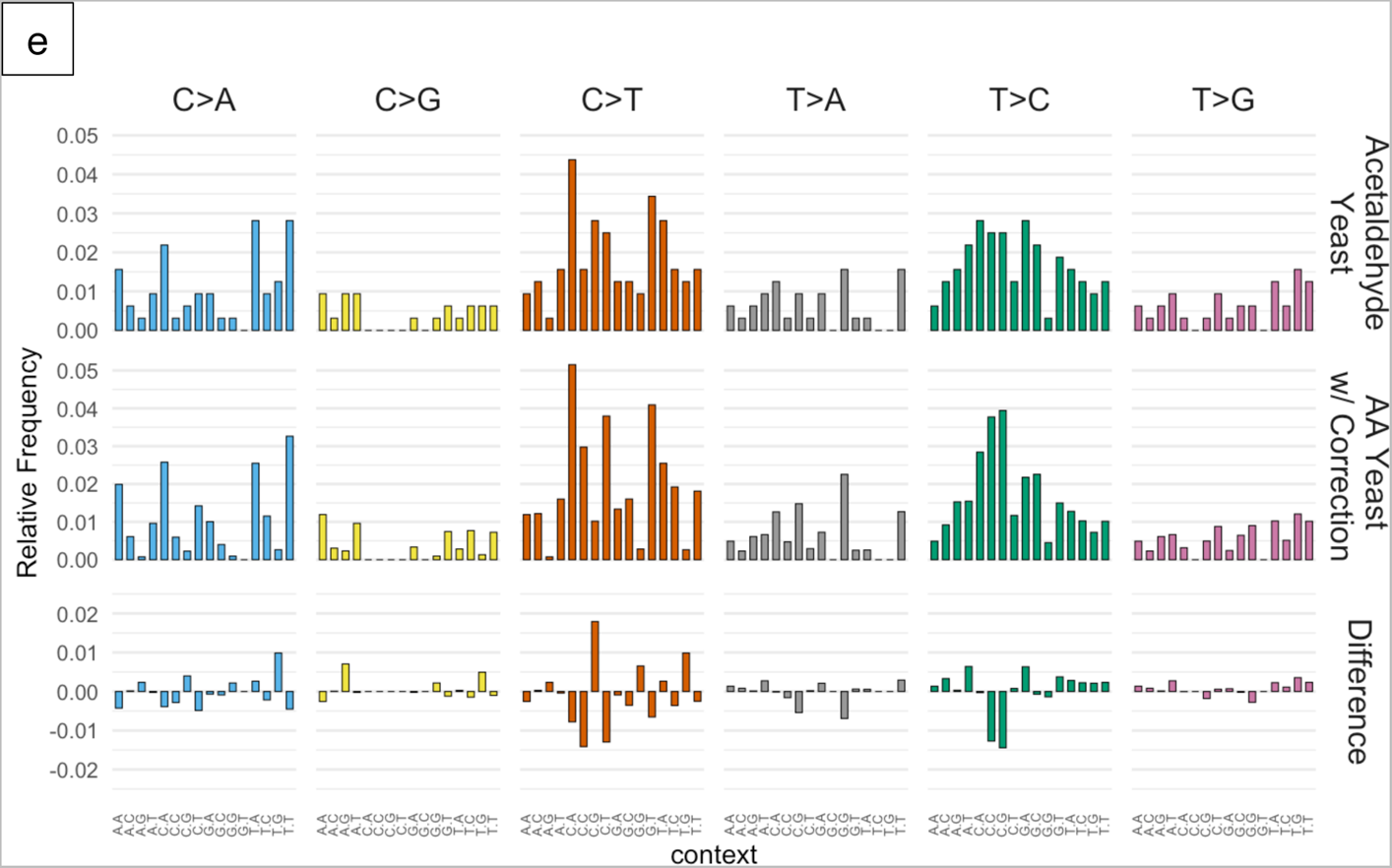
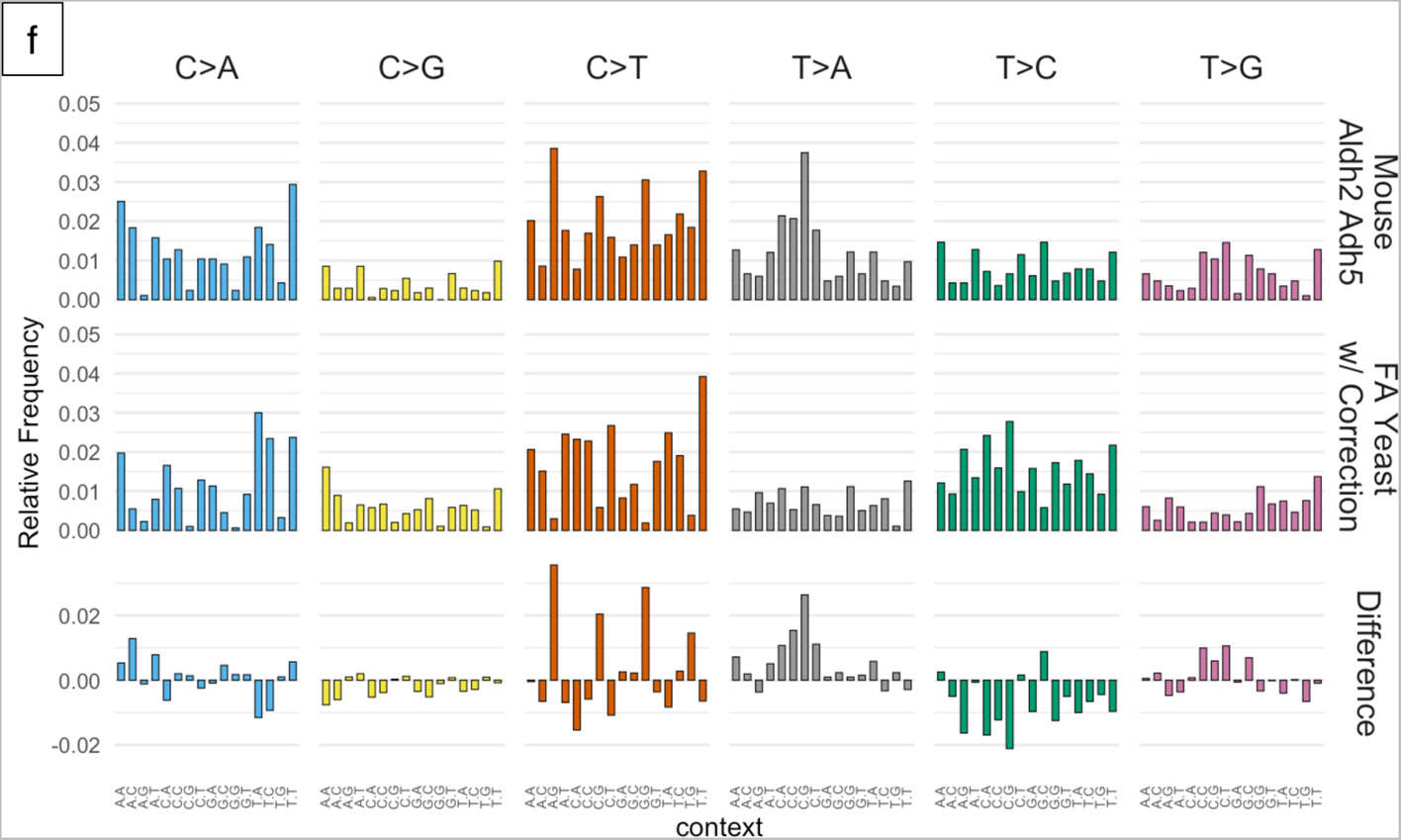
Comparisons of mutational patterns between (a) untreated controls and formaldehyde; (b) untreated controls and acetaldehyde; (c) formaldehyde and acetaldehyde; (d) formaldehyde with and without correction for trinucleotide frequencies in mouse; (e) acetaldehyde with and without correction for trinucleotide frequencies in mouse; and (f) Aldh2 Adh5 deficient mouse cells and trinucleotide frequency corrected yeast treated with formaldehyde.

A recent high profile paper described mutational patterns obtained in mice that were genetically deleted for genes important in aldehyde detoxification, ADH5 and ALDH2, thus leading to buildup of endogenous aldehydes (37). To compare our mutational patterns derived from mutagenized yeast genomes to these profiles from mice, we first adjusted for differences in trinucleotide abundances between the two species to obtained corrected mutational patterns (see Figure 4d and 4e). A main difference between the yeast and mouse genomes is the lower abundance of <u>C</u>pG motifs in the latter. Nonetheless, the corrected mutational patterns retained high similarity to the original (uncorrected) patterns in yeast (cosine similarity values > 0.95).

When we compared the various mutational patterns, we noticed that cosine similarity values are relatively low when comparing between the corrected yeast patterns we derived and the mouse patterns from Dingler et al. (37) (see Table 1). These values are somewhat higher when comparing the formaldehyde pattern in yeast to the mouse patterns. A closer examination of these profiles suggests that there are likely to be mutations from other sources mixed in the mouse patterns, e.g. from SBS1 (deamination of 5-methylcytosine at CpG motifs, see Figure 4f). We also noted some differences among the various mouse patterns themselves: while the ones from Adh5^−/-^ and Aldh2^−/-^ Adh5^−/-^ had cosine similarity = 0.887, the Aldh2^−/-^ profile was noticeably more dissimilar (cosine similarity < 0.8 vs. the other two profiles). This is consistent with the likelihood that there are other mutation sources mixed in with the mouse mutational patterns, which may be confounding interpretation of a hypothesized pattern induced by excess endogenous aldehydes.

**Table 1:** Cosine similarity values between mutational profiles of (FA) formaldehyde- and (AA) acetaldehyde-mutagenized yeast (with correction for trinucleotide frequencies in mouse) and of mice deficient for aldehyde detoxification genes from (37).

|  | Mouse Aldh2<sup>-/-</sup> | Mouse Adh5<sup>-/-</sup> | Mouse Aldh2<sup>-/-</sup><br>Adh5<sup>-/-</sup> |
| --- | --- | --- | --- |
| Yeast FA, corrected | 0.658 | 0.735 | 0.767 |
| Yeast AA,<br>corrected | 0.617 | 0.633 | 0.673 |

**Table 2:**
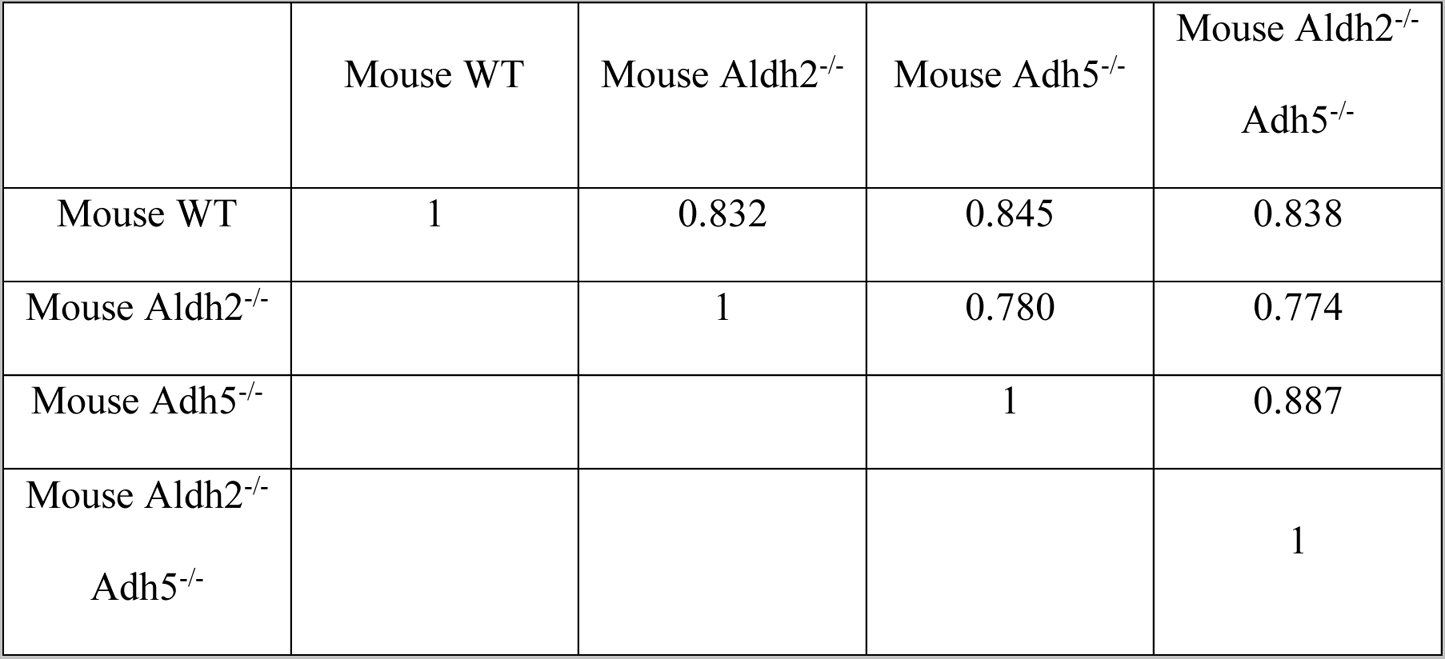
Cosine similarity values among mice deficient for aldehyde detoxification genes from (37).

|  | Mouse WT | Mouse Aldh2<sup>-/-</sup> | Mouse Adh5<sup>-/-</sup> | Mouse Aldh2<sup>-/-</sup><br>Adh5<sup>-/-</sup> |
| --- | --- | --- | --- | --- |
| Mouse WT | 1 | 0.832 | 0.845 | 0.838 |
| Mouse Aldh2<sup>-/-</sup> |  | 1 | 0.780 | 0.774 |
| Mouse Adh5<sup>-/-</sup> |  |  | 1 | 0.887 |
| Mouse Aldh2<sup>-/-</sup><br>Adh5<sup>-/-</sup> |  |  |  | 1 |

### Formaldehyde mutational pattern resembles COSMIC SBS signature 40

We then investigated whether these mutational patterns might shed light on the etiology of any known COSMIC mutational signatures. We started with the mouse profiles published by Dingler et al. (37) and confirmed that none of the mouse profiles showed a particularly close resemblance to any known COSMIC signature (see Figure 5a). All of those cosine similarity values were < 0.8, suggesting that if there are bona fide COSMIC signatures within the mouse mutational patterns, they are possibly obscured by being in a mixture of multiple signatures.

**Figure 5:**
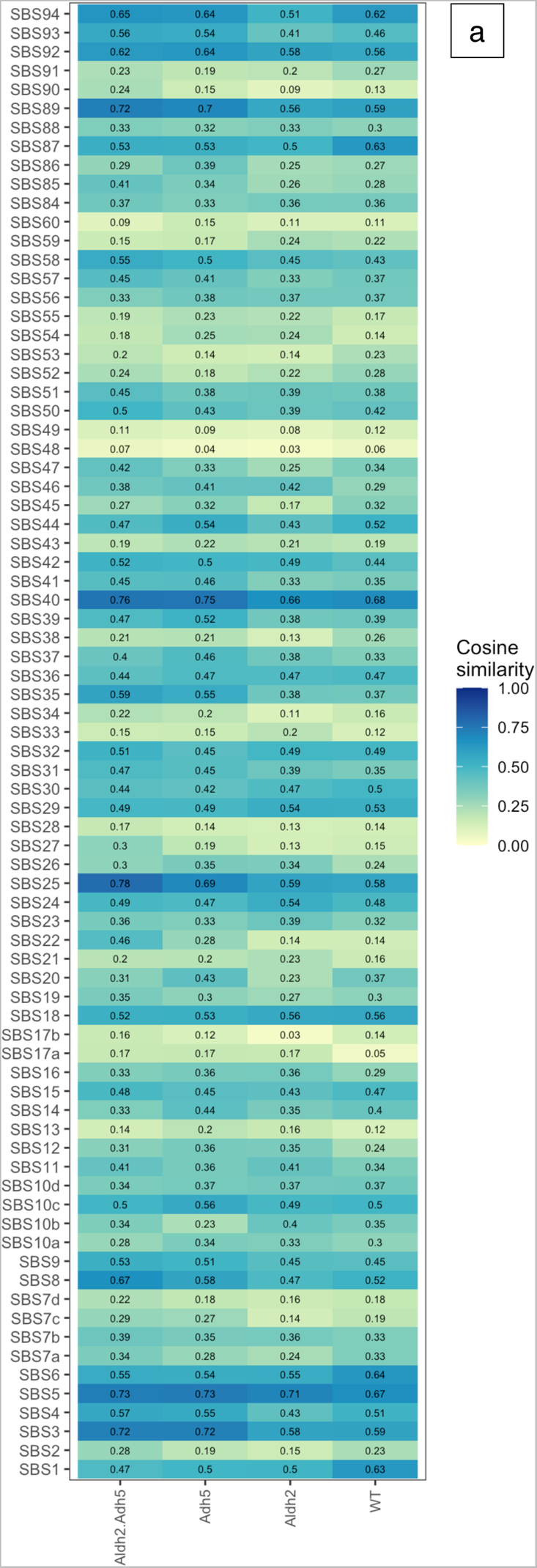
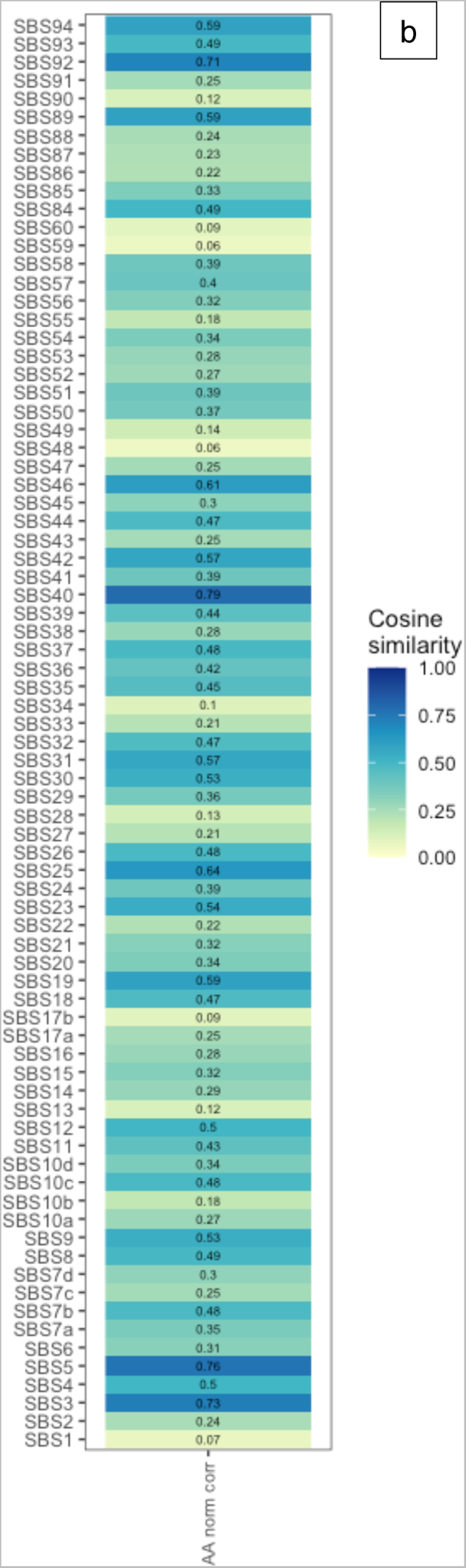
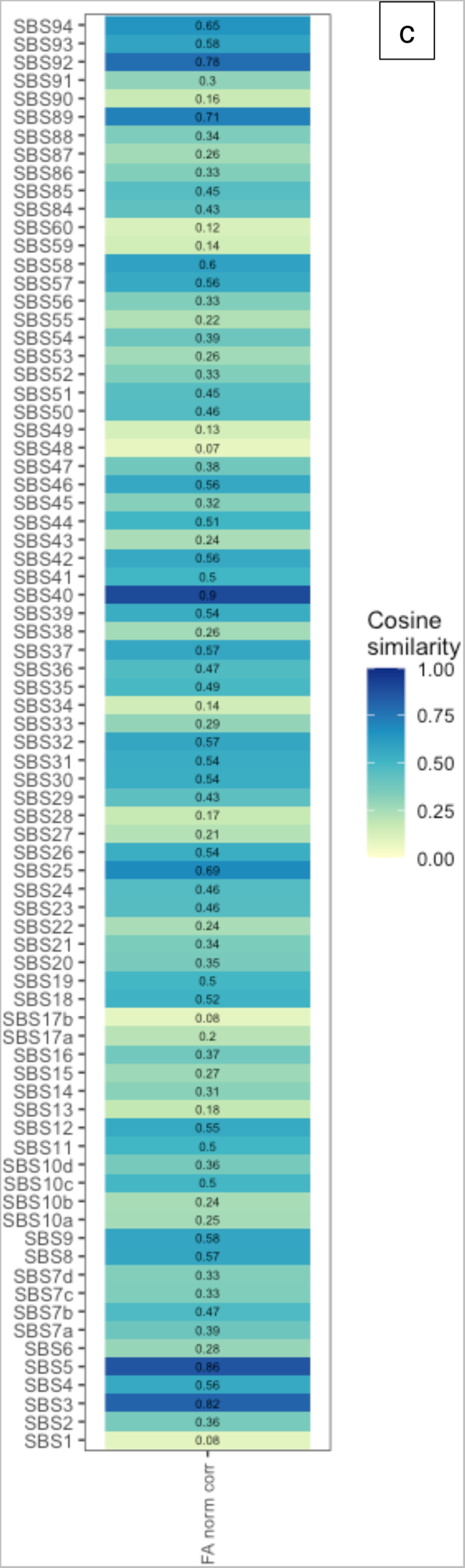
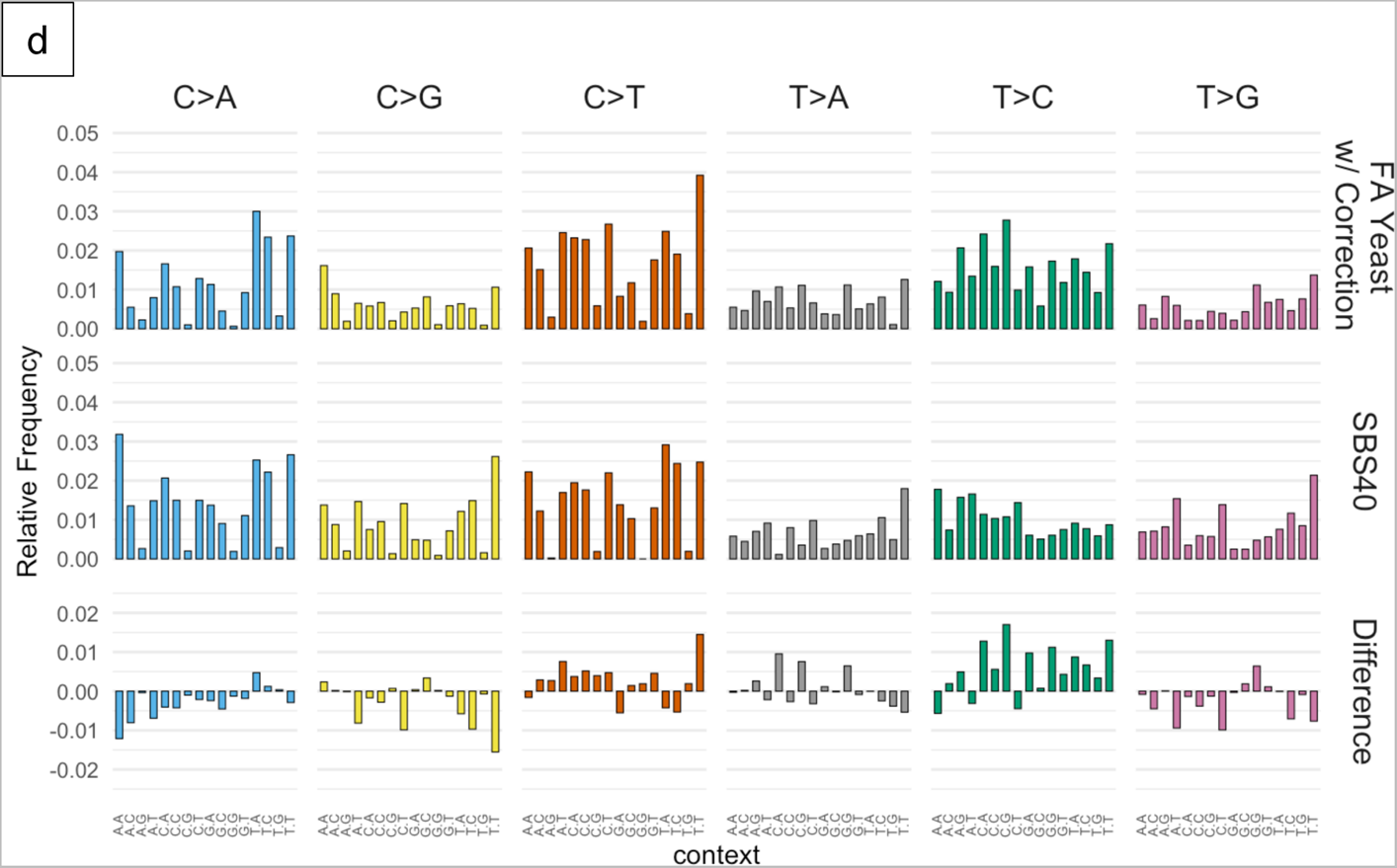
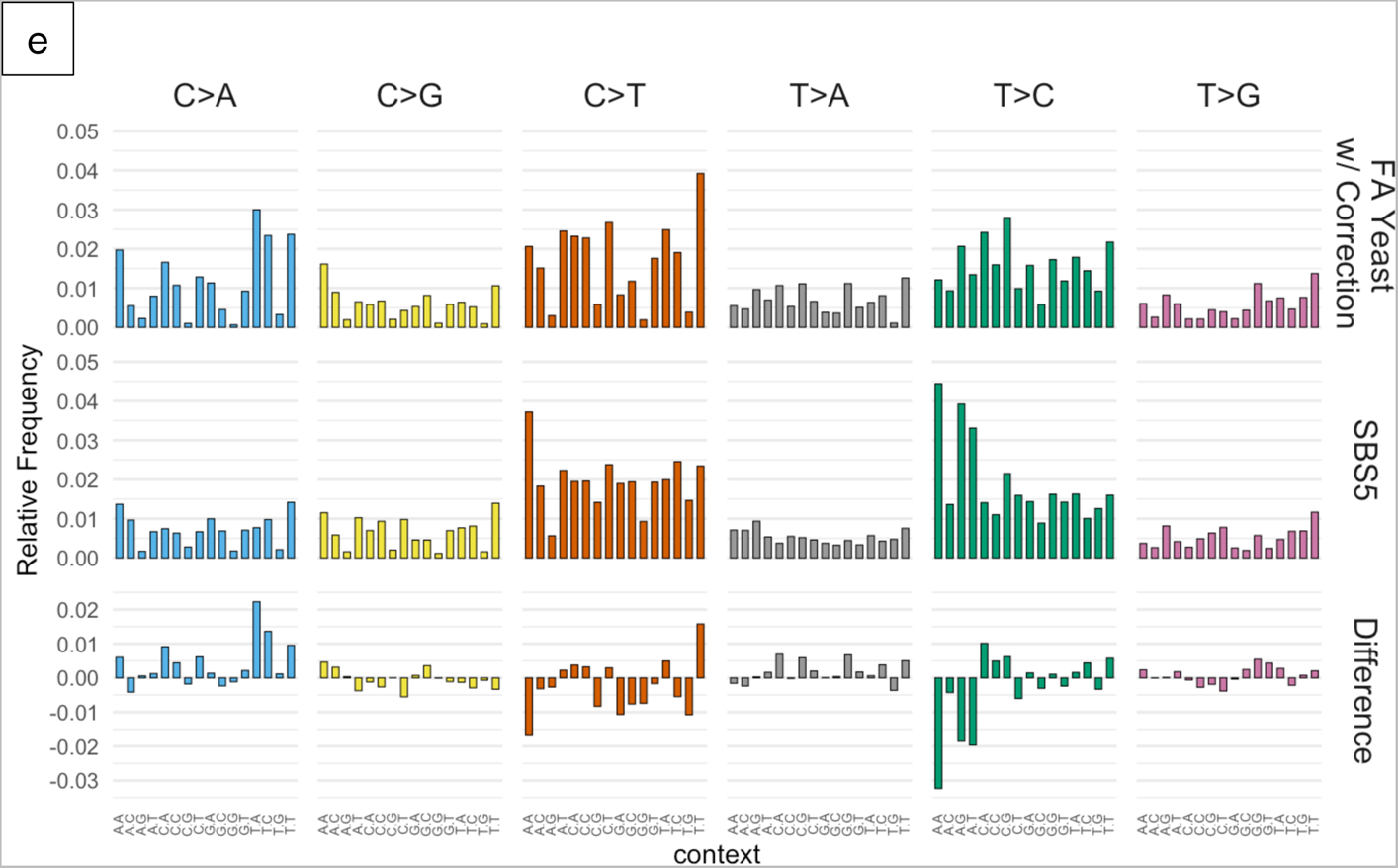
(a) Cosine similarity values comparing mouse mutational patterns from Dingler et al. vs. COSMIC SBS signatures. (b) Cosine similarity values comparing acetaldehyde mutational pattern in yeast (with trinucleotide abundance correction) vs. COSMIC SBS signatures. (c) Cosine similarity values comparing corrected formaldehyde mutational pattern in yeast vs. COSMIC SBS signatures. (d) Comparison of corrected formaldehyde mutational pattern in yeast vs. SBS signature 40. (e) Comparison of corrected formaldehyde mutational pattern in yeast vs. SBS signature 5.

Comparison of the corrected acetaldehyde pattern from yeast vs. known COSMIC signatures also yielded, at best, cosine similarity of 0.79 to SBS40, a signature of unknown etiology (see Figure 5b). Since we had better direct control of the induced mutagenesis experiments using the yeast system with exogenously applied mutagen, it does not seem as likely that other mutagenic processes are obscuring the acetaldehyde-induced pattern. We conclude that the acetaldehyde pattern we obtained is not a plausible match for any known COSMIC signature at this point.

Finally, we compared the formaldehyde pattern to the COSMIC signatures, finding that the closest match is to SBS40, with cosine similarity = 0.9 (see Figures 5c and 5d). The second closest match was to SBS5 (cosine similarity = 0.864, see Figures 5c and 5e). We previously studied an SBS5-like mutational pattern in yeast and showed that similar patterns are widely conserved in many species. Moreover, the SBS5-like pattern is due to error-prone translesion DNA synthesis in the absence of added mutagens and increases with increasing sugar metabolism (53). An SBS40-like mutational pattern would require a separate explanation, which would be the addition of exogenous formaldehyde to our experimental system. As such, we propose that a plausible etiology for SBS40 in cancers is the mutagenicity of formaldehyde.

## Discussion

In this paper, we report the use of a sensitive ssDNA-based mutagenesis reporter system to characterize the mutagenic properties of two small aldehydes, formaldehyde and acetaldehyde. This system is especially well suited for investigating chemical agents with relatively weak mutagenicity. A challenge of using conventional mutagenesis systems to study weak mutagens is that induced mutations can be rare and it can be difficult to discern a reliable mutational pattern using relatively few mutations (36,37).

In addition to being a more sensitive reporter system, it is also considerably more cost-efficient to sequence compact yeast genomes (each ∼12 megabases) than mammalian genomes which are much larger (∼3 gigabases). By applying a correction to account for different abundances of trinucleotide motifs, we can use data from the sequencing of mutagenized yeast to infer the expected mutational pattern in another species. Another key advantage is the single-stranded configuration of the DNA precludes repair that requires a complementary strand. By sidestepping intervention from DNA repair processes, the ssDNA system can provide, in effect, a purer readout of the effects of mutagenesis per se. Leveraging these advantages of the ssDNA mutagenesis reporter system, we were able to infer the mutational patterns of both formaldehyde and acetaldehyde.

The two small aldehydes share some similar mutagenicity characteristics, but also have their differences. Both induce dose-dependent increases in mutagenesis at lower concentrations. But whereas formaldehyde-induced mutagenesis essentially plateaus from 4 mM up to 8 mM, acetaldehyde-induced mutagenesis peaks at 75 mM and then drops sharply at the even higher concentration of 100 mM. Both aldehydes induce significant cytotoxicity at the higher end of their respective ranges of tested concentrations, but yeast are able to tolerate considerably higher doses of acetaldehyde overall. Yeast are presumably evolved to cope with significantly higher concentrations of acetaldehyde, since it is an abundant intermediate in ethanol production from fermentation (54). Both aldehydes cause an excess of C/G > A/T transversions, which is consistent with previous reports showing preferential adduct formation and mutagenesis at guanines (55–61). Interestingly, acetaldehyde induces an excess of deletion variants of five or more bases in our system, but formaldehyde does not, consistent with previous reports (55,62). These various mutagenic characteristics of formaldehyde and acetaldehyde reflect their chemical similarities and differences.

The 96-channel mutational patterns of formaldehyde and acetaldehyde revealed further differences between the two compounds. Whereas the acetaldehyde pattern did not particularly resemble any known COSMIC signature, new mutational signatures will be revealed as more cancer samples are sequenced and analyzed. Since alcohol consumption is associated with multiple cancer types and it is thought that the acetaldehyde from alcohol detoxification would surely damage DNA (63), associated mutational signature(s) may yet be discovered in the future. The formaldehyde pattern we obtained was similar to SBS signature 40. SBS40 is currently of unknown etiology, but it is known to be present in at least 28 cancer types (28), making SBS40 the third most common mutational signature in cancers. The high prevalence of SBS40 hints at an endogenous origin for the underlying DNA damage that is present in different cell types throughout the body. Since formaldehyde is produced endogenously and exists at steady state concentrations in humans in the range of tens of micromolar (33), it would fit this profile. When all of the available information is taken into consideration, mutagenesis from endogenously generated formaldehyde emerges as a plausible candidate for the etiology of SBS40.

Comparison with mutational patterns from mice deleted for aldehyde detoxification genes suggest that those profiles are likely mixtures of mutations from different mutagenic processes, and not just from DNA damage due to accumulation of excess endogenous aldehydes. For example, the contribution from SBS1 (C/G > T/A at <u>C</u>pG motifs) was quite noticeable. Mutagenesis from other sources likely interferes with making an accurate inference of the aldehyde-associated mutagenesis. This is perhaps another significant challenge when using systems for mutational detection that are not (and maybe can not) be properly controlled to factor out mutagenesis from other sources. Deployment of more specialized and sensitive mutagenesis detection systems where the experimenters have more direct control over the mutation induction can continue to play an important role in shining new light on mutagenesis.

## Acknowledgments

We thank S. Gelova, B. Xhialli, and N. Liang for their critical feedback on this work. The authors gratefully acknowledge support from a Tier 2 Canada Research Chair, an NSERC Discovery Grant, an Ontario Early Researcher Award, and uOttawa startup funding to K.C. Author contributions are as follows: All authors performed experiments, and analyzed/discussed the data; K.C. conceived and supervised the study, and wrote the draft manuscript. The authors declare there are no competing interests.

## Notes

### Competing Interest Statement

The authors have declared no competing interest.

